# ResistanceGA: An R package for the optimization of resistance surfaces using genetic algorithms

**DOI:** 10.1101/007575

**Authors:** William E. Peterman

**Affiliations:** Illinois Natural History Survey, Prairie Research Institute, University of Illinois, 1816 S. Oak Street, Champaign, IL 61820 USA

**Keywords:** CIRCUITSCAPE, circuit theory, cost distance, gene flow, genetic algorithm, landscape genetics, least cost path, resistance distance, resistance optimization, resistance surface

## Abstract

1. Understanding how landscape features affect functional connectivity among populations is a cornerstone of landscape genetic analyses. However, parameterization of resistance surfaces that best describe connectivity is largely a subjective process that explores a limited parameter space.
2. ResistanceGA is a new R package that utilizes a genetic algorithm to optimize resistance surfaces based on pairwise genetic distances and either CIRCUITSCAPE resistance distances or cost distances calculated along least cost paths. Functions in this package allow for the optimization of both categorical and continuous resistance surfaces, as well as the simultaneous optimization of multiple resistance surfaces.
3. There is considerable controversy concerning the use of Mantel tests to accurately relate pairwise genetic distances with resistance distances. Optimization in uses linear mixed effects models with the maximum likelihood population effects parameterization to determine AICc, which is the fitness function for the genetic algorithm.
4. ResistanceGA fills a void in the landscape genetic toolbox, allowing for unbiased optimization of resistance surfaces and for the simultaneous optimization of multiple resistance surfaces to create a novel composite resistance surface.

## Introduction

First coined in 2003, landscape genetics has experienced rapid growth in both the number of studies and range of analytical methods utilized (Manel et al. 2003; Storfer et al. 2010). This integrative field draws on landscape ecology, spatial statistics, and population genetics to address a wide range of questions. One of the primary goals of landscape genetic studies has been to understand how landscape features affect spatial genetic structure (Manel *et al.* 2003; Storfer et al. 2007). Rather than simply assessing isolation-by-distance, many landscape genetic studies quantify the effective distance (i.e. resistance distance) between spatially distinct sample locations as a function of the landscape matrix (Spear et al. 2005; McRae 2006). In the absence of direct observation of movement or dispersal across the landscape, resistance distances are often interpreted as functional connectivity (e.g., Cushman et al. 2006). Critically, to infer functional connectivity and resistance distance, an appropriate resistance surface must be parameterized. As defined by Spear et al. (2010), a resistance surface is a spatial layer that assigns a value to each landscape or environmental feature, with values representing the extent to which that feature impedes or facilitates connectivity for an organism.

Resistance values of surfaces have been determined using a variety of methods, including: habitat suitability models (e.g., Wang et al. 2008), telemetry (e.g., Driezen et al. 2007), and most commonly expert opinion (Murray et al. 2009). Less often, parameterization is informed by empirical movement studies (e.g., Stevens et al. 2006) or spatial prediction of ecological processes (Peterman et al. 2014). All of these are acceptable approaches, but each comes with its own caveats. Of particular concern is the fact that expert opinion often fails to accurately describe the biological or ecological process(es) being modeled (Shirk et al. 2010; Charney 2012). Even when biological or ecological processes are known and explicitly modeled, there is no guarantee that these processes will relate meaningfully to the movement of genes across the landscape (Peterman et al. 2014).

Ultimately, the assignment and evaluation of resistance values generally covers only a limited parameter space and remains a trial and error process to determine the best resistance parameters. Graves, Beier and Royle (2013) attempted to alleviate the subjectivity of resistance surface parameterization by using a search algorithm to maximize Mantel /-correlation between inter-individual genetic distance and least cost path distance. While their optimization procedures recovered the maximum Mantel */* when it existed, they found that resistance estimates were often imprecise and much smaller than simulated resistance values. They also found that response surfaces were quite flat, making identification of a global optimum difficult. In contrast, Peterman *et al.* (2014) found well-defined global optima when using Ricker and monomolecular data transformations in combination with optimization algorithms. However, the optimization procedure utilized by Peterman et al. (2014) is limited to continuous resistance surfaces (e.g., temperature, moisture, percent canopy cover) and requires an inefficient search of all possible data transformations.

There are numerous challenges to optimizing resistance surfaces based on pairwise (genetic) distance data. Among these challenges is the fact that resistance surfaces can have a high dimensionality, as in land use, land cover surfaces. Another issue is that there currently is no closed-form expression to determine the landscape resistance values that describe pairwise distances, potentially making optimization intractable with gradient-based algorithms. Finally, landscape features and environmental gradients do not exist in isolation. Therefore, an ideal solution to resistance surface optimization will be to simultaneously optimize multiple resistance surfaces to create a composite resistance surface. The ResistanceGA package for the R programming environment (R Core Team 2014) has been developed to address these issues, and to fill a void in the landscape genetic toolbox. While the initial impetus for this package was landscape genetic analyses, but any pairwise measures across the landscape (e.g., movement rates) could be utilized to optimize resistance surfaces.

## Description

### Genetic algorithms

Genetic algorithms (GAs) represent a powerful and flexible stochastic optimization framework for finding solutions to both discrete and continuous optimization problems (Holland 1975). Inspired by biological principles, genetic algorithms create a population of individuals (offspring) with traits (parameters to be optimized) encoded on “chromosomes”. The genotypes (parameter combinations) of each individual solve the fitness function, and the fittest individuals from each generation survive to reproduce (Sivanandam & Deepa 2007). The GA evolution process is facilitated by exploration and exploitation (Scrucca 2013). Exploration of parameter space occurs through random generation of new parameter values resulting from mutation, as well as exchange of genetic information through crossover. Exploitation ultimately reduces diversity in the population by selecting the fittest individuals each generation. The population continues to evolve until a sufficient number of generations have passed without an improvement in fitness (Scrucca 2013).

### Resistance optimization

ResistanceGA utilizes the general-purpose genetic algorithm from the GA R package (ga function; Scrucca 2013). Briefly, the optimization proceeds as follows:

1. The original raster surface is imported into R. If the surface is continuous, it is rescaled to range of 0-10, preserving the relative spacing between all levels.
2. The evolution process starts by generating a random initial population of size *n*. If a continuous surface, the selected parameters determine (a) which of eight transformations will be applied (Fig. 1); (b) the shape of the transformation; (c) the maximum value of the transformation. If a categorical surface, each level of the resistance surface is reclassified to the values of the parameters.
3. Using the spatial locations where genetic samples have been collected, either CIRCUITSCAPE (McRae *et al.* 2008; McRae & Shah 2009) is called from R to calculate pairwise resistance distances across the landscape created in step 2 above, or cost distances are calculated from least cost paths using the package (van Etten 2014).
4. A linear mixed effects model with a maximum likelihood population effects parameterization (MLPE) is fit to the data. Pairwise genetic distance is the response and the scaled and centered pairwise resistance distance is the predictor. The MLPE mixed effects parameterization is used to account for the non-independence among the pairwise data (Clarke, Rothery & Raybould 2002).
5. The Akaike information criterion (AIC) is obtained from the fitted MLPE model. AICc is then calculated by penalizing the model for the number of parameters used during transformation/reclassification. AICc is the fitness measure that the genetic algorithm seeks to maximize. Because the ga function works through maximization, the sign of AICc is reversed.
6. Steps 2-5 are repeated until the specified number of *n* individuals have been created. The genetic algorithm then conducts selection on the population, and the individuals with the best AICc are carried over to the next generation to form the reproducing population. A new population is then formed through mutation and crossover.
7. Steps 2-6 are repeated until the specified number of generations have passed without improvement to the AlCc.

**Figure 1.**
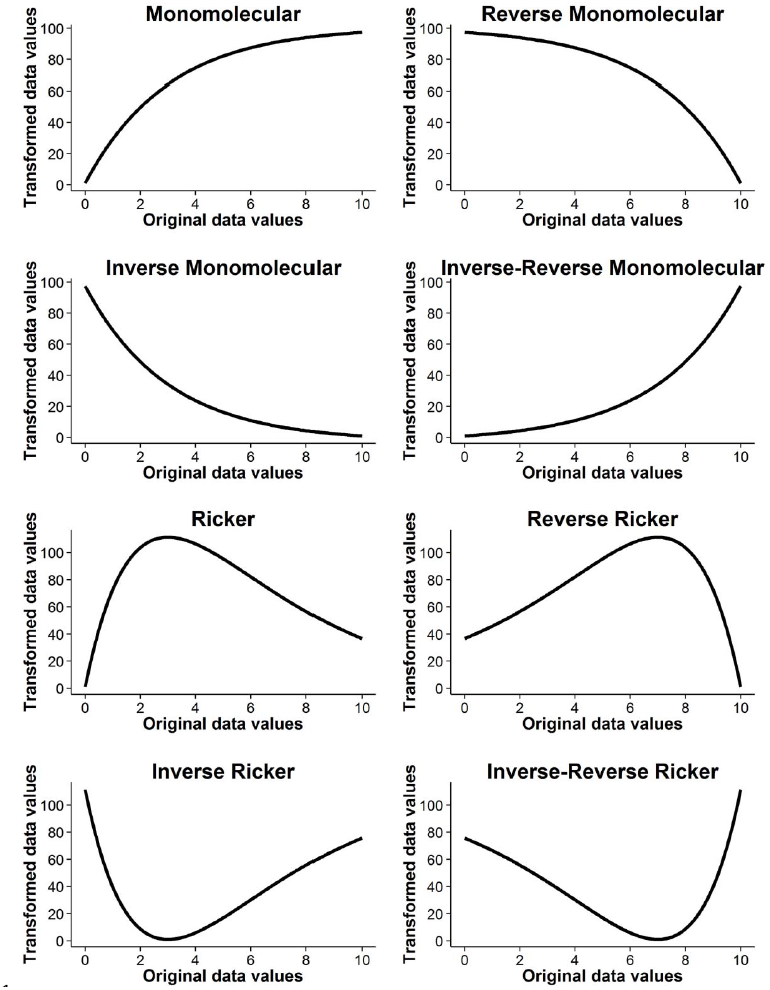
There are eight continuous resistance surface data transformations implemented in ResistanceGA. Prior to transformation, the original continuous resistance surface had values ranging from 0-10. The shape and magnitude of each transformation are each controlled by a single parameter. All transformations in the figure have a shape parameter value of 3, and maximum value parameter of 100. Linear relationships are not explicitly incorporated, but all monomolecular functions become linear as the shape parameter increases.

### Continuous surfaces

There are eight transformations that can be applied to continuous surfaces (Fig. 1). Each transformation relies on either the Ricker or monomolecular functions (Bolker 2008), as well as rescaling functions to keep values in positive parameter space. The shape and magnitude of each transformation are controlled by two parameters, which are adjusted during optimization (Peterman et al. 2014). During optimization, the genetic algorithm searches different combinations of transformations, scale parameters, and shape parameters. Linear transformations are not explicitly included, but all of the monomolecular functions become linear as the shape parameter increases in value. In this way, linear responses can effectively be modeled without increasing the number of transformations for the genetic algorithm to test.

### Categorical surfaces

Categorical or feature surfaces, such as land cover or roads, can also be optimized using ResistanceGA. A surface is considered categorical in ResistanceGA if it contains 15 or fewer levels. To make this process tractable, it is necessary to hold the value of one level constant throughout optimization. Because pairwise resistance values are relative, failing to hold one level constant can result in multiple equivalent solutions, and the algorithm may fail to reach an optimal solution (e.g., relative resistance values of 1, 5, and 10 are equivalent to 2, 10, and 20).

Combining resistance surfaces Resistance surfaces are simultaneously optimized by sequentially modifying each surface to be optimized. Continuous and categorical surfaces are each modified as described above, then all modified surfaces are summed together to create a single, composite resistance surface, which is then used to calculate pairwise resistance distances or cost distances.

## Overview of ResistanceGA functions

The optimization functions passed to ga rely heavily on the R package raster (Hijmans 2014) to import, export and manipulate spatial raster (*.asc*) files, as well as lme4 (Bates et al. 2014) to fit mixed effects models. To visualize continuous surface transformations, ResistanceGA uses the graphing features of ggplot2 (Wickham 2009). The functions available in ResistanceGA are summarized in Table 1.

**Table 1:** Functions of the ResistanceGA package

| **Function** | **Returned objects** | **Description** |
| --- | --- | --- |
| Combine_Surfaces | R raster object | Combine multiple resistance surfaces into a new composite surface based on specified parameters |
| CS.prep | Named list | This function prepares and bundles the inputs necessary to run CIRCUITSCAPE |
| Diagnostic.Plots | .tif file | This function returns a multipanel figure including the histogram of residuals and qqplot from a fitted mixed effects model |
| GA.prep | Named list | This function prepares and compiles the objects and commands needed to execute the genetic algorithm |
| gdist.prep | Named list | This function prepares and bundles the inputs necessary to run gdistance |
| Grid.Search | contour plot, R data object | For a single continuous surface, this function can be used to visualize the AICc response surface |
| lower | R data object | Convenience function to obtain the lower half of a square distance matrix |
| MLPE.lmm | lmer object | Fits a maximum likelihood population effects mixed effects model in lme4 |
| MS_optim | GA object, diagnostic plots, .txt summary file | This is a wrapper function for simultaneously optimizing multiple surfaces. |
| Plot.trans | ggplot object | Implements and plots a continuous surface transformation |
| Resistance.opt_multi | AICc value | This is the function passed to ga to optimize multiple resistance surfaces simultaneously |
| Resistance.opt_single | AICc value | This is the function passed to ga to optimize resistance surfaces in isolation |
| Resistance.tran | R raster object, .asc file | Applies specified transformation to continuous resistance surface and returns a raster object. Optionally, a .asc file can be exported |
| Run_CS | R raster object or R data object | This function executes CIRCUITSCAPE from R and returns either the lower half of the pairwise resistance distance matrix or the cumulative current map produced by CIRCUITSCAPE |
| Run_gdistance | costDistance matrix object from gdistance | This function executes gdistance and returns the costDistance matrix object |
| SS_optim | .tif files, .csv summary tables | This is a wrapper function for optimizing surfaces in isolation. All surfaces in a common directory will be optimized in turn, and numerous summary tables of optimized parameters and AICc values are produced |

## Implementation

Resistance surfaces can be optimized using cost distances (least cost path) or using circuit-based resistance distances. Cost distances are calculated using the R package gdistance (van Etten 2014), while electrical current resistances are calculated using the free, open-source software CIRCUITSCAPE (v4.0 or greater; McRae & Shah 2009). If using CIRCUITSCAPE, input data formats must conform to those of CIRCUITSCAPE (McRae & Shah 2009). Currently, CIRCUITSCAPE can only be executed from R through ResistanceGA on computers using Windows operating systems. Prior to running any of the optimization functions (MS_optim, SS_optim, Resistance.Opt_multi or Resistance.Opt_single) the CS.prep/gdist.prep, and GA.prep preparation functions must be run. These functions create and format the inputs and data objects necessary to run dRCUITSCAPE/gdistance, parameterize the genetic algorithm, and fit MLPE mixed effects models. These functions have been developed to require a minimum input from the user. For instance, at minimum, all that needs to be specified to run GA. prep is the directory where the *.asc* files to be optimized reside. However, all available arguments of the ga function can be set by the user to modify the genetic algorithm. The arguments and default settings of the preparation functions are described in Table 2.

**Table 2:**
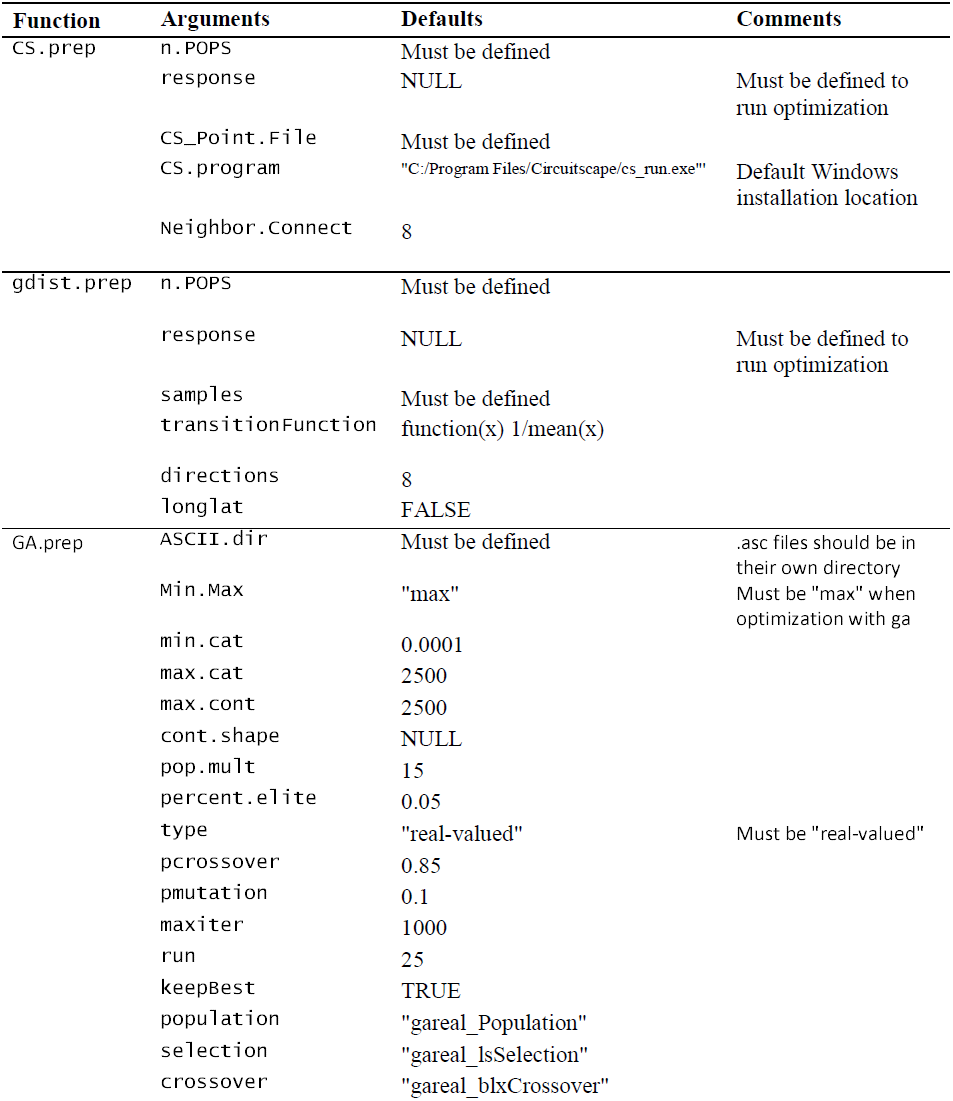
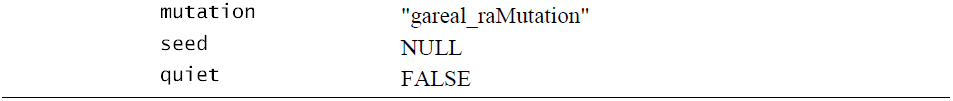
Arguments of the required preparation functions and their default settings. Only one of CS.prep or gdist.prep needs to be run, depending upon whether optimization will use CIRCUITSCAPE or gdistance.

There is no established framework for optimization of resistance surfaces, but an envisioned work flow is detailed in Figure 2. Genetic algorithms are stochastic optimization procedures; therefore it is highly advised to run all optimizations at least twice to confirm convergence and parameter estimates. Also, because boundaries are placed on the parameter space searched, if the optimized resistance values are at or near the limits set, the optimization should be rerun after expanding the search space. It is important to note that simultaneous optimization of multiple surfaces often results in parameter values that differ from truth. However, the relative resistance values of the resultant resistance surface are highly or perfectly correlated with truth (see worked example), and the resistance surfaces are identical once they are rescaled to a minimum resistance of one. Users of ResistanceGA should be aware of this fact, and interpret results accordingly.

**Figure 2.**
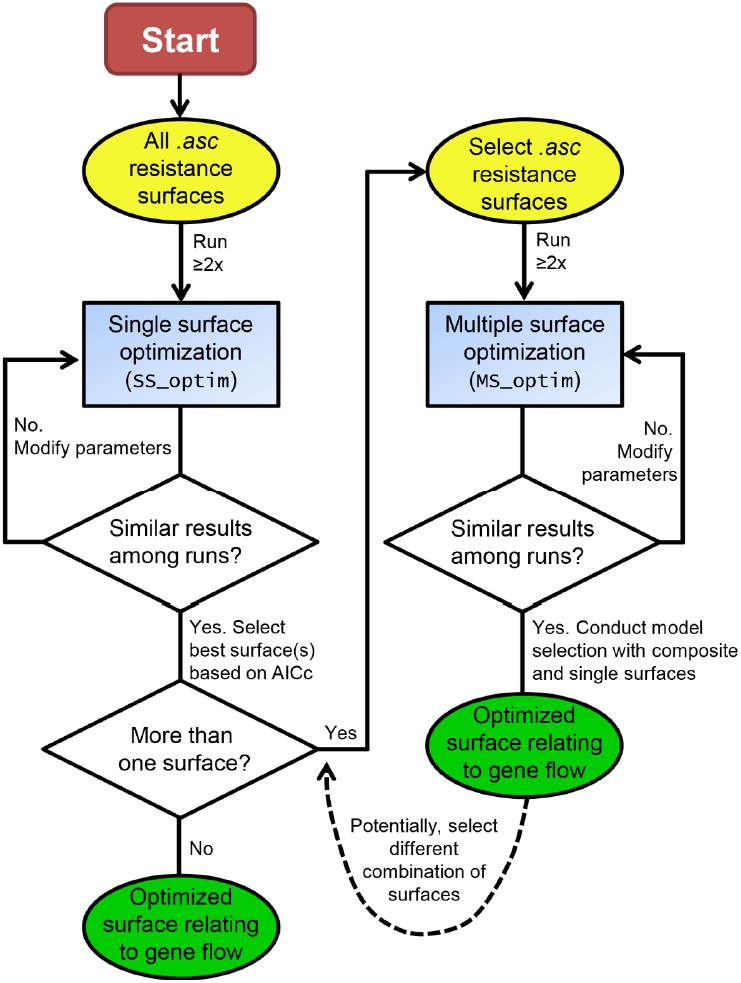
Flow chart depicting a potential work flow for optimizing resistance surfaces. Analysis begins by optimizing each resistance surface in isolation. If multiple surfaces are supported (e.g., ΔAICc < 4), these surfaces can be simultaneously optimized to create a composite surface. If desired, different combinations of the best supported single surfaces can be optimized and compared. Inputs into the optimization are shown in yellow ovals, optimization steps are blue rectangles, decision points are white diamonds, and final products from optimization are green ovals.

Genetic algorithms are effective at finding an optimal solution, but the can be computationally intensive. To ensure that parameter space is adequately searched, the population of individuals produced each generation must be of sufficient size. In ResistanceGA the default setting is to produce a population that is 15 times the number of parameters being optimized, up to a maximum population size of 100 individuals. For example, when a continuous surface is being optimized, 45 individuals (3 parameters x 15) will be produced each generation, and a typical optimization takes 50-300 generations. This results in the creation of 2 250-13 500 resistance surfaces and dRCUITSCAPE/gdistance runs. Therefore, the greatest impediment to optimizing resistance surfaces is time. Both the spatial extent and the number of sample locations in the analysis affect the time necessary to calculate pairwise resistance or cost distances. On a computer with an Intel i7 3.4 GHz processor, 25 sample locations distributed across a 250 x 250 cell resistance surface takes ∼5 seconds to complete an iteration with CIRCUITSCAPE. The same surface with 100 sample locations takes ∼14 seconds. Finally, 25 sample locations on and a 1 000 x 1 000 cell surface can take up to 90 seconds to complete. Because CIRCUITSCAPE calculates resistance over all possible pathways, it is more computationally intensive than calculating least cost paths. On average, optimization with gdistance is about three times faster than calculating resistance distances with CIRCUITSCAPE. To further reduce the optimization time when using least cost paths, the optimization can be run in parellel by setting parallel = TRUE in GA.prep. Additional strategies to reduce the runtime of CrRCUITSCAPE/gdistance include modifying the connection scheme (Neighbor.Connect/directions) from the default setting of 8 to 4, and reducing the resolution of the resistance surfaces (i.e. increase cell size). The latter will reduce the total number of cells on the surface, and generally has limited effect on the quantification of relative resistances across the landscape (McRae et al. 2008).

## Worked examples

The examples below use simulated data provided with ResistanceGA. Code to generate the example resistance surfaces and spatial point data can be found in Appendix S1. More extensive examples and further descriptions of the functions included in ResistanceGA can be found in the vignette included with the package.

### Example 1: Single surface optimization

This example optimizes a single resistance surface using CIRCUITSCAPE.

~~~
# Get data

# Install ‘devtools’ package, if needed if(!(“devtools” %in% list.files(.libPaths()))) {
 install.packages(“devtools”)

}

library(devtools)
install_github(“wpeterman/ResistanceGA”) # Download package
library(ResistanceGA) # Load  package
rm(list = ls())

#  Create directory  for  reading/writing CIRCUITSCAPE files and results
if(“ResistanceGA_Examples”%in%dir(“C:/”)==FALSE)
 dir.create(file.path(“C:/”, “ResistanceGA_Examples”))

# Create a subdirectory for the first example
dir.create(file.path(“C:/ ResistanceGA_Examples/”,“SingleSurface”))

# Directory to write .asc files and results

write.dir <- “C:/ ResistanceGA_Examples/SingleSurface/”

# Load example data, write continuous surface as ‘.asc’ file
data(resistance_surfaces)

continuous <- resistance_surfaces[[2]]

writeRaster(continuous,filename=paste0(write.dir,“cont.asc”),overwrite=TRUE)

# Load sample location data
data(samples)
write.table(samples,
            file=paste0(write.dir,“samples.txt”),
            sep=“\t”,col.names=F,row.names=F)

# Create a spatial points object for plotting
sample.locales <- SpatialPoints(samples[,c(2,3)])

# Run preparation  functions

GA.inputs <- GA.prep(ASCII.dir=write.dir,

                 max.cat=500,
                 max.cont=500,
                 seed=555,
                 quiet=TRUE)

CS.inputs <- CS.prep(n.POPS=length(sample.locales),
                 CS_Point.File=paste0(write.dir,“samples.txt”),
                 CS.program=‘“C:/Program   Files/Circuitscape/cs_run.exe”’)

# Transform the continuous surface

r.tran <- Resistance.tran(transformation=“Monomolecular”,
                  shape=2,
                  max=275,
                  r=continuous)

# Visualize  the transformation (Fig. 3)
plot.t <- Plot.trans(PARM=c(2,275),

                 Resistance=continuous,
                 transformation=“Monomolecular”)

# Calculate pairwise  resistance values  to use as  the  response
CS.response <- Run_CS(CS.inputs=CS.inputs,
                 GA.inputs=GA.inputs,
                 r=r.tran)

# Rerun ‘CS.prep’ to include response
CS.inputs <- CS.prep(n.POPS=length(sample.locales),
                 response=CS.response,
                 CS_Point.File=paste0(write.dir,“samples.txt”),
                 CS.program=‘“C:/Program   Files/Circuitscape/cs_run.exe”‘)

# Run optimization
SS_RESULTS <- SS_optim(CS.inputs=CS.inputs,
                         GA.inputs=GA.inputs)

# Compare results
SS_table <- data.frame(c(“Monomolecular”, 2.0, 275),
                     t(SS_RESULTS$ContinuousResults[c(3:5)]))
colnames(SS_table)  <-  c(“Truth”, “Optimized”)
SS_table
                  Truth         Optimized
                  Equation Monomolecular  Monomolecular

                  shape    2      1.999999
                  max      275      274.9982
~~~

The exact transformation parameters were recovered in 147 iterations (Fig 3).

**Figure 3.**
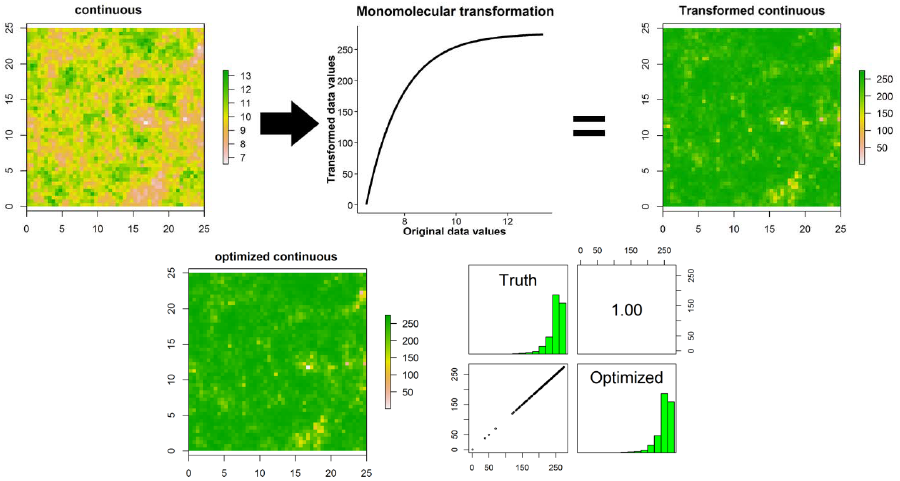
Optimization of a continuous surface that was transformed using a monomolecular function following Example 1 in the main text (transformation = Monomolecular, shape = 2.0, maximum = 275). Optimization with CIRCUITSCAPE took 147 iterations of the genetic algorithm and precisely recovered the transformation parameters resulting in perfectly correlated resistance surfaces.

### Example 2: Multisurface optimization

This example simultaneously optimizes three resistance surface using least cost paths.

~~~
# Get data
data(resistance_surfaces)
data(samples)

# Create a spatial points object
sample.locales <- SpatialPoints(samples[,c(2,3)])

# Create a subdirectory for the second example
dir.create(file.path(“C:/ResistanceGA/”,“MultipleSurfaces”))

# Run ‘GA.prep’
GA.inputs <- GA.prep(ASCII.dir=resistance_surfaces,

                 Results.dir=“C:/ResistanceGA/MultipleSurfaces/”,
                 max.cat=500,
                 max.cont=500,
                 seed = 321,
                 parallel = 4) # Run on in parallel on 4 cores

# Run ‘CS.prep’ functions
gdist.inputs<-gdist.prep(n.POPS=length(sample.locales),

                 samples=sample.locales)

# Set parameters for transforming and combining  resistance surfaces
PARM <- c(1,250,75,6,3.5,150,1,350)

#  PARM<- c(1,    # First feature of categorical

#       250,    # Second feature of categorical

#       75,     # Third feature  of categorical

#       6,      # Transformation equation for continuous surface

#       3.5,    # Shape parameter

#       150,    # Scale parameter

#       1,      # First feature of feature surface

#       350)    # Second feature of feature surface

# Combine resistance surfaces
Resist <- Combine_Surfaces(PARM=PARM,
                gdist.inputs=gdist.inputs,
                GA.inputs=GA.inputs,
                out=NULL,
                rescale=TRUE)

# Create the response surface
gd.response <- Run_gdistance(gdist.inputs=gdist.inputs,
                                GA.inputs=GA.inputs,
                                r=Resist)

# Re-run ‘gdist.prep’ to include the response
gdist.inputs<-gdist.prep(n.POPS=length(sample.locales),

                        response=lower(as.matrix(gd.response)),
                        samples=sample.locales)

# Run ‘MS_optim’
Multi.Surface_optim.gd <-  MS_optim(gdist.inputs=gdist.inputs,
                                       GA.inputs=GA.inputs)

# View optimized parameter values with the true simulation values
Summary.table <- data.frame(PARM,round(t(Multi.Surface_optim.gd@solution),2))
colnames(Summary.table)<-c(“Truth”, “Optimized”)
row.names(Summary.table)<-c(“Category1”, “Category2”, “Category3”,
“Transformation”, “Shape”, “Max”, “Feature1”, “Feature2”)
Summary.table
                 Truth Optimized
Category1          1.0      1.00
Category2         250.0    268.27
Category3          75.0     80.42
Transformation      6.0      6.51
Shape               3.5      3.50
Max               150.0    161.11
Feature1            1.0      1.00
Feature2          350.0    375.56

# Compare the true and optimized resistance surfaces
optim.resist <-  Combine_Surfaces(PARM=Multi.Surface_optim.gd@solution,

                  gdist.inputs = gdist.inputs,
                  GA.inputs = GA.inputs)

resist.stack <-  stack(Resist,optim.resist)
names(resist.stack) <- c(“Truth”, “Optimized”)
pairs(resist.stack) # Correlation  plot (Fig. 4)
plot(resist.stack) # Resistance maps (Fig. 4)
~~~

This example took 210 iterations to optimize. As described above, the optimization has recovered different parameter values than those that were simulated, but the relative values of the simulated and optimized surfaces are identical (Fig. 4).

**Figure 4.**
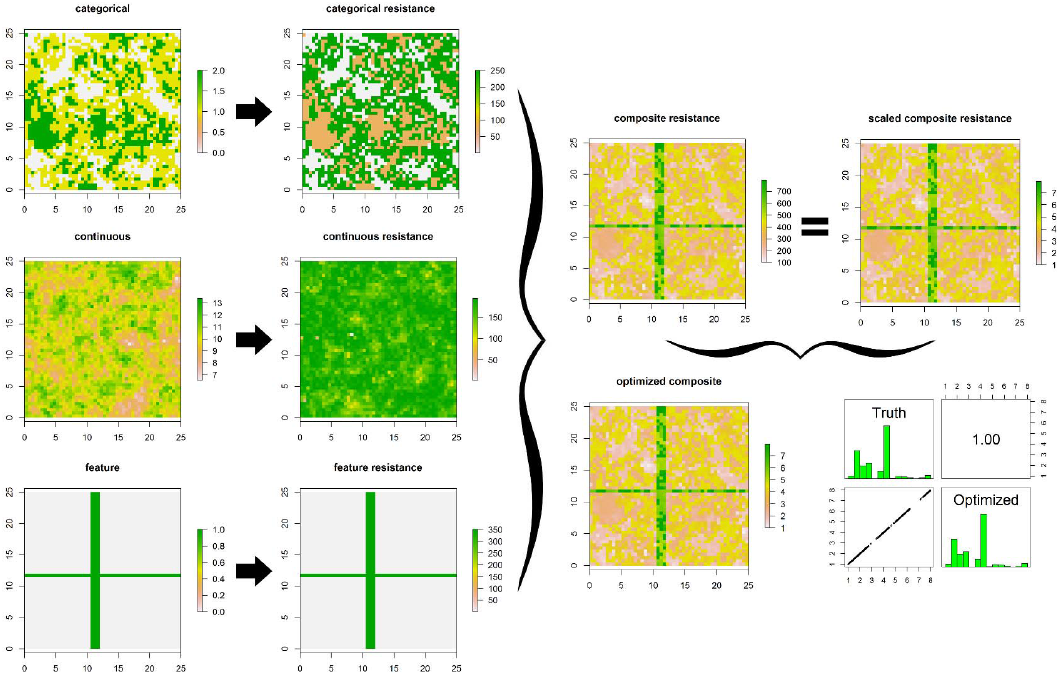
Simultaneous optimization of three resistance surfaces (Example 2 in the main text). The original categorical, continuous, and feature resistance surfaces are shown on the left and their actual resistances following transformation/rescaling are shown in the middle. When these surfaces are added together to create a composite, the minimum resistance value is no longer 1, so the surface is rescaled to present the relative resistance values (scaled composite resistance). Optimization using least cost paths took 215 iterations of the genetic algorithm when run in parallel on 4 cores. The relative resistance values of the optimized surface and the true resistance surface are perfectly correlated.

## Obtaining ResistanceGA

ResistanceGA is hosted on GitHub, and can be downloaded using the install.github function from the package:

> install.github(“wpeterman/ResistanceGA”)

Following download, the package can be loaded:

> library(ResistanceGA)

View the vignette with more demonstrations of ResistanceGA functions:

> vignette(‘ResistanceGA’)

Alternatively, the HTML vignette can be viewed here: http://goo.gl/n7VOZx

## Acknowledgements

I thank G. Connette for many discussions concerning the development and implementation of functions in ResistanceGA, and for comments on an early draft of this manuscript.

## Data accessibility

Example data are provided with ResistanceGA or can be made using the code provided in Appendix S1.

## Supporting information

Appendix S1. R code to generate the example resistance surfaces provided with ResistanceGA

